# Circular single-stranded DNA is a superior homology-directed repair donor template for efficient genome engineering

**DOI:** 10.1101/2022.12.01.518578

**Authors:** Keqiang Xie, Jakob Starzyk, Ishita Majumdar, Katerina Rincones, Thao Tran, Danna Lee, Sarah Niemi, John Famiglietti, Bernhard Suter, Richard Shan, Hao Wu

**Affiliations:** Full Circles Therapeutics, Inc. 625 Mt. Auburn St., Ste. 211, Cambridge, MA 02138; Stellate DNA, Inc. 625 Mt. Auburn St., Ste. 211, Cambridge, MA 02138

**Keywords:** circular single-stranded DNA, genome editing, CRISPR/Cas9, homology-directed repair, CAR-T

## Abstract

The toolbox for genome editing in basic research and therapeutic applications is rapidly expanding. While efficient targeted gene ablation using nuclease editors has been demonstrated from bench to bedside, precise transgene integration remains a technical challenge. AAV6 has been a prevalent donor carrier for homology-directed repair (HDR) mediated genome engineering but has reported safety issues, manufacturing constraints, and restricted applications due to its 4.5 Kb packaging limit. Non-viral targeted genetic knock-ins rely primarily on double-stranded DNA (dsDNA) and linear single-stranded DNA (lssDNA) donors. Both dsDNA and lssDNA have been previously demonstrated to have low efficiency and cytotoxicity. Here, we developed a non-viral <u>g</u>enome writing <u>catalyst</u> (GATALYST™) system which allows production of ultrapure, mini<u>c</u>ircle <u>s</u>ingle-<u>s</u>tranded <u>DNAs</u> (cssDNAs) up to ∼20 Kb as donor templates for highly efficient precision transgene integration. cssDNA donors enable knock-in efficiency of up to 70% in induced pluripotent stem cells (iPSCs), superior efficiency in multiple clinically relevant primary cell types, and at multiple genomic loci implicated for clinical applications with various nuclease editor systems. When applied to immune cell engineering, cssDNA engineered CAR-T cells exhibit more potent and durable anti-tumor efficacy than those engineered from AAV6 viral vectors. The exceptional precision and efficiency, improved safety, payload flexibility, and scalable manufacturability of cssDNA unlocks the full potential of genome engineering with broad applications in therapeutic development, disease modeling and other research areas.

**Highlights:**

- Scalable production of mini<u>c</u>ircle <u>ssDNA</u> (cssDNA) with highly engineered phagemid system
- <u>G</u>enome writing <u>catalyst</u> (GATALYST™) system with cssDNA donor template demonstrates superior efficiency and safety in various cell types and genomic loci
- GATALYST gene writing system enables ultra-large transgene integration
- cssDNA engineered CAR-T outperforms AAV engineered CAR-T with superior anti-tumor function

## Introduction

The RNA-guided Cas9 nucleases from the microbial CRISPR (clustered regularly interspaced short palindromic repeat)-Cas systems (CRISPR-Cas9) are robust and versatile tools for targeted genome editing in eukaryotic cells (Cho *et al*., 2013; Cong *et al*., 2013; Jinek *et al*., 2013; Mali *et al*., 2013). CRISPR-Cas9 offers unprecedented opportunities to modify genome sequences in primary human cells throughout the field of cell and gene therapies. Cas nucleases are directed to a region in the genome by a guide RNA (gRNA) and induce a targeted double-strand break (DSB), where the resulting cellular repair mechanisms can be exploited to induce either error-prone or defined alterations. In the absence of a repair template, the cell typically repairs the lesion by a non-templated repair pathway such as nonhomologous end joining (NHEJ). In the presence of a homology-directed repair (HDR) DNA template, precise repair directed by regions of homology on DNA donor templates is triggered.

Previously, large disease modifying transgenes were introduced into various cell types via lentivirus or retrovirus. These methods harbored inherent challenges, which led to technical challenges, resulting in undesirable phenotypes like cancer. Such challenges include hard to track insertion genomic locus and copy number, difficulty manufacturing quality viral particles, and intrinsic heterogeneity of engineered cells. Targeted transgene insertion has been successfully demonstrated in the chimeric antigen receptor (CAR) T cell field using DNA donor template encapsulated in recombinant associated Adenovirus (rAAV) (Gaj *et al*., 2017). However, in addition to the potential viral-related safety and manufacturing challenges, the donor template payload application of rAAV is restricted by the packaging limit of 4.5 Kb.

Non-viral targeted genome editing, especially large transgene insertion, represents a significant unmet need, technically and medically, in the field of immune oncology and monogenic disorders. Non-viral targeted gene knock-in usually relies on double-strand DNA (dsDNA) donors, either in linear or circular form, to achieve targeted DNA insertion at a locus of interest (Salsman & Dellaire, 2017). However, the use of dsDNA suffers from low efficiency and high cytotoxicity. More recently, single-stranded DNAs (ssDNAs) have been proven more effective than dsDNAs templates as donors for HDR in CRISPR-based genome editing due to reduced cellular toxicity and increased HDR efficiency (*Bai et al., 2020*; Iyer *et al*., 2022; Li, 2019; Miura *et al*., 2018; Quadros *et al*., 2017; Roth *et al*., 2018; Shy *et al*., 2022).

Moreover, illegitimate random integration is expected to be less frequent for ssDNA compared to dsDNA templates (Won & Dawid, 2017; Wurtele *et al*., 2003; Zorin *et al*., 2005). ssDNA has become a groundbreaking tool for a wide variety of applications including engineering of human primary T cells (Shy *et al*., 2022). An endogenous T cell receptor (TCR) replaced with NY-ESO-1 antigen-specific 1G4 TCR encoded in ∼2 Kb long ssDNA demonstrated superior therapeutic efficacy compared to traditional methods involving TCR insertion using lentivirus (Roth *et al*. 2018). In addition, a hybrid ssDNA HDR template encoding an anti B-cell maturation antigen (BCMA) CAR targeting the human T-cell receptor (TCR) alpha chain (*TRAC*) locus in primary T cells revealed superior cellular immunophenotype and rapid *in vivo* tumor clearance (Shy *et al*., 2022). The ssDNA donors used in these seminal studies were exclusively linear molecules produced with *in vitro* methods (Li, 2019; Shy *et al*., 2022). They were produced by manipulation of DNA with polymerase chain reaction (PCR) coupled with enzymatic degradation of one of two DNA strands. However, the enzymatic production of linear ssDNA (lssDNA) templates is inherently inefficient and only commercially available at up to 5 Kb, due to accumulation of mutations from polymerase chain reaction, high reagent cost, and low scalability, thus limiting successful clinical application.

More recently, phagemid-derived long circular ssDNA (cssDNA) HDR donors were demonstrated to have higher efficiency of HDR-based repair in multiple cell lines (HEK293T and K562 cells) (Huh, 2019; Iyer *et al*., 2022). The cssDNA HDR templates outperform other forms of DNA donor templates due to its increased HDR efficiency, high specificity, large length capacity, and low cytotoxicity.

Here, we developed a technology platform using cssDNA purified from engineered phagemids for targeted gene writing. We established that cssDNA donor templates enable higher knock-in efficiency with decreased cytotoxicity when compared to lssDNA or dsDNA templates in cell lines, iPSC cells, and clinically relevant primary cells, demonstrating its potential application for genetic disorders and immune cell therapy.

## Results

### Production of cssDNA in a Highly Engineered Phagemid System

We have developed a proprietary M13 phagemid system that allows us to efficiently produce cssDNA. After co-transformation with a helper plasmid in XL1-blue *E. coli*, the single strand of the phagemid vector is packaged into phage particles and secreted into the culture media (**Figure 1A**). The endotoxin-free cssDNA phage particles were then isolated using anion exchange chromatography column. Over 100 cssDNAs ranging from 1 Kb to 22 Kb in lengths have been purified using this engineered phagemid system. Agarose gel imaging showed over 95 % cssDNA purity across a range of lengths (**Figure 1B**). No negative correlation between the yield and the length of the cssDNA molecules up to 22 Kb (**Figure 1C**), thus, the yield of cssDNA is not expected to be compromised for larger cssDNA. A varying yield was observed across different cssDNA constructs, though. Moreover, the cssDNA yield from multiple production batches for the same cssDNA molecule was consistent (**Figure 1D**), suggesting the production process is reproducible and easily scalable. ssDNA structures are highly flexible, and their functional form is thermodynamically less stable. The circular form of ssDNA derived from phagemid is considered more resistant to exonuclease than lssDNA. Our purified cssDNA was very stable and did not degrade after 14 days of storage at room temperature and withstood up to 42 freeze/thaw cycles over this period (**Supplemental Figure 1**). To determine the stability of cssDNA in a cellular context, a qPCR method was developed to quantify the amount of dsDNA or cssDNA from the total DNA extracted from cell lysate. Once dsDNA or cssDNA was delivered into mammalian K562 cells by electroporation, cell pellets were collected at different time points and the amount of dsDNA or cssDNA were quantified by qPCR. The half-life of the circular phagemid dsDNA in the cells was much longer than cssDNA with half-life 42 hours vs. 5.8 hours (**Figure 1E**). This demonstrates that cssDNA is indeed less stable than dsDNA. However, the shorter half-life of cssDNA in cells could be advantageous for genome editing to minimize the undesired cellular toxicity and recombination.

**Figure 1.**
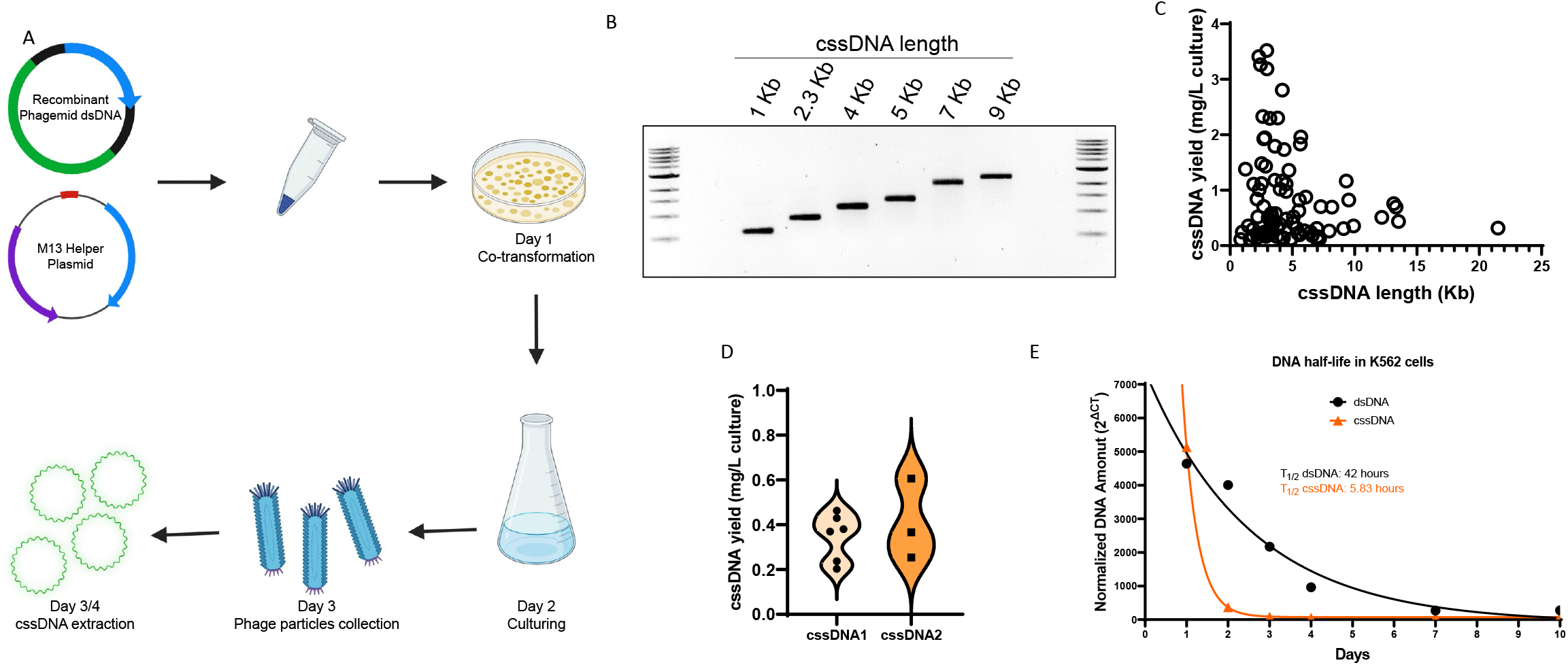
Scalable cssDNA manufacturing and purification from bacteriophage. **A**. Schematic diagram of cssDNA manufacturing. Recombinant phagemid containing sequence of interest was co-transformed with M13 helper plasmid into *E. coli*. Phage particles were produced and collected from culture supernatant. cssDNA was then extracted and purified from phage. **B**. Representative DNA agarose gel images showing the purity of manufactured cssDNA at lengths ranging from 1 Kb to 9 Kb. **C**. cssDNA yields (mg per liter of culture volume) for ∼100 of cssDNA molecules at various lengths. **D**. The violin plot of the production of same cssDNA molecules over different batches. **E**. cssDNA and dsDNA half-life analysis in K562 cells. cssDNA or dsDNA were electroporated into K562 cells. The remaining DNAs in cells were analyzed by quantitative PCR. DNA was normalized with 2^ΔCT^ methods. Half-life (T_1/2_) were obtained after one phase decay non-linear curve fitting.

### GATALYST Gene Writing Platform with cssDNA Outperforms Other Non-Viral Nucleic Acid Payloads

To directly compare the knock-in efficiency with different types of donor template, GFP reporter DNA donor templates flanked with ∼300 nt 5’ homology arm and ∼300 nt 3’ homology arm targeting the *RAB11A* locus in the form of double-stranded circular plasmid, linear single-stranded or circular single-stranded DNA were tested in K562 cells. As illustrated in **Figure 2A**, when DNA donor templates were co-electroporated with ribonucleoprotein (RNP) complex consisting of spCas9 protein and single guide RNA (sgRNA) targeting exon 1 of the *RAB11A* locus (Roth *et al*., 2018), precise HDR of the locus was evaluated by GFP reporter expression using flow cytometry. Day 7 post electroporation, knock-in efficiency, as measured by GFP^+^ cell percentage, was much higher with cssDNA as the repair donor template, reaching to over 40% with 3 µg of cssDNA (**Figure 2B**). In contrast, reduced knock-in efficiency was observed with lssDNA or dsDNA donor templates. As expected, cell health was not compromised by cssDNA or lssDNA; however, the cell viability of 3 µg dsDNA engineered cells was less than 10% (**Figure 2C**). Higher knock-in efficiency and lower cell toxicity suggests cssDNA is a superior donor DNA template over lssDNA and dsDNA for HDR-mediated genome knock-in. To examine this further, we then tested the cssDNA knock-in efficiency at various timepoints post electroporation across a range of dosages. **Figure 2D** showed that cssDNA-mediated knock-in at the *RAB11A* locus plateaued on Day 7 and was sustained for at least 2 weeks. When knock-in efficiency (examined on Day 7) was plotted together with cell viability (examined on Day 3 post electroporation), a clear dose-dependent increase of knock-in efficiency for up to 5 µg of cssDNA emerged, all while maintaining more than 80% cell viability (**Figure 2E**). When higher concentrations of cssDNA (to 14 µg) were used, both knock-in efficiency and cell viability were reduced. Therefore, maximal knock-in efficiency is achieved with ∼5 µg cssDNA. cssDNA donor templates targeting *RAB11A* locus with different lengths of the 5’ and 3’ homology arms were also tested (**Supplementary Figure 2A)**. Knock-in efficiency data indicated that 300 nt homology arm of each end resulted in the highest HDR efficiency, and there is no additional benefit when increasing the lengths up to 1500 nt (**Supplementary Figure 2B)**. Thus, HDR cssDNA donor templates with 300 nt homology arms lengths were used for the following genome engineering studies.

**Figure 2.**
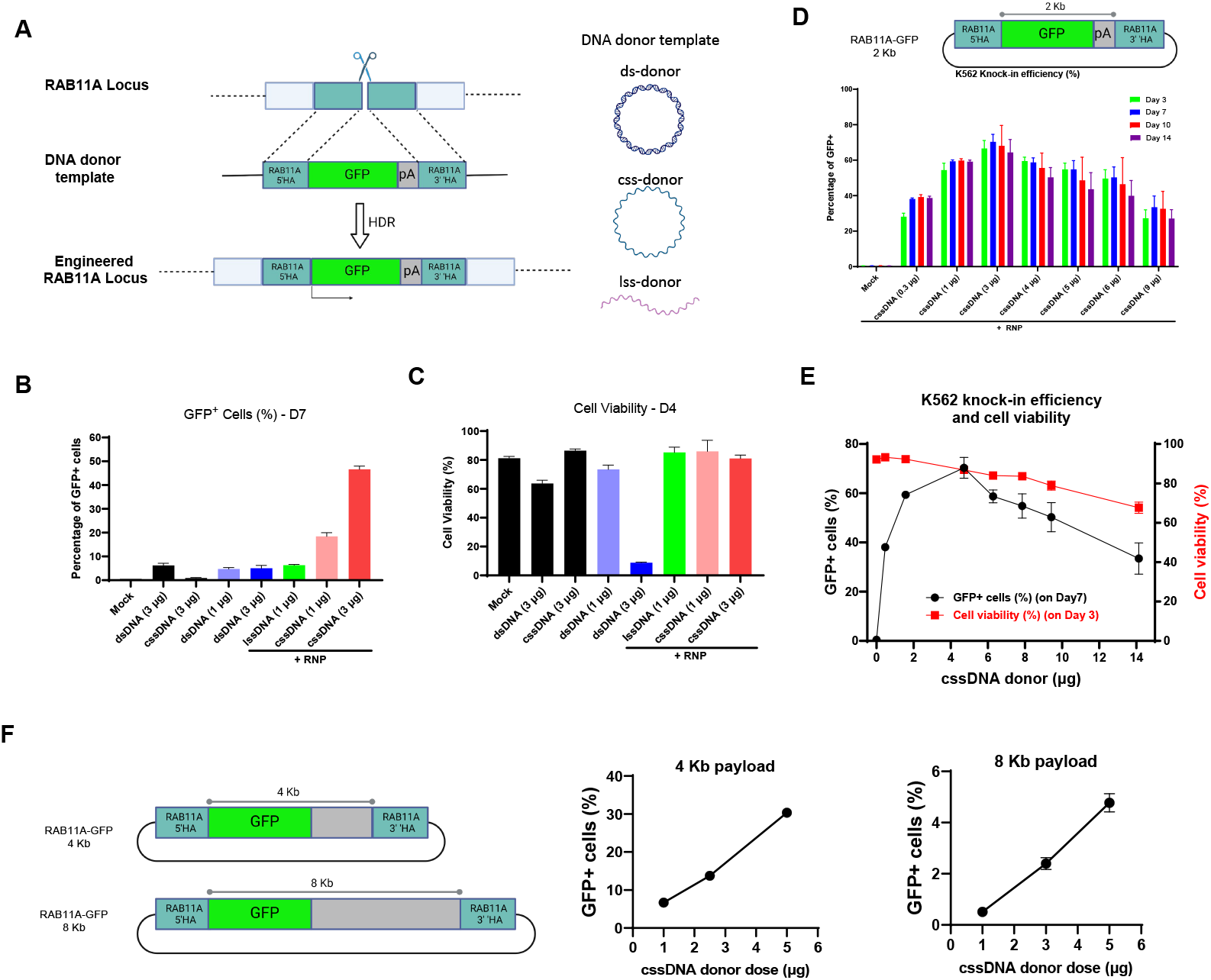
GFP reporter knock-in efficiency and cell viability in K562 cells. **A**. Schematic diagram of the CRISPR-mediated genomic integration of a GFP reporter gene at the *RAB11A* locus with dsDNA, cssDNA or lssDNA donor templates. **B**. The GFP reporter knock-in efficiency at the *RAB11A* locus in K562 cells after engineering with cssDNA, lssDNA or dsDNA as donor templates on Day 7 post electroporation. **C**. Cell viability of K562 cells on Day 4 after engineering with cssDNA, lssDNA or dsDNA as donor templates. **D**. Knock-in efficiency of GFP reporter gene (2 Kb payload) at the *RAB11A* locus in K562 cells over time with increasing amounts of cssDNA donor template. **E**. Knock-in efficiency and cell viability with increasing amount of cssDNA donor template in K562 cells. 2 Kb GFP reporter payload cssDNA targeting the *RAB11A* locus was used to engineer K562 cells. Knock-in efficiency was measured on Day 7 and cell viability was assessed on Day 4 after electroporation. **F**. Schematic diagram of the 4Kb and 8Kb GFP reporter payload cssDNAs targeting *RAB11A* locus (l*eft*). Knock-in efficiency (measured on Day 7) for the 4 Kb payload (*middle*) and 8 Kb payload (*right*) with increasing donor template concentrations.

We further tested the knock-in efficiency at the *RAB11A* locus in K562 cells for larger (4 Kb and 8 Kb) payload sizes using cssDNA (**Figure 2F**). Although lower than the 2 Kb payload, both the 4 Kb and 8 Kb payload cssDNAs produced significant knock-in efficiency in a dose-dependent manner (**Figure 2F**). Therefore, the GATALYST gene writing platform with cssDNA outperformed other nucleic acids templates for gene knock-in and enabled efficient knock-in of large (up to 8 Kb) transgenes.

### Highly Efficient iPSC Engineering with cssDNA Donor Template

Genetic engineering, especially precise transgene knock-in of induced pluripotent stem cells (iPSCs), holds great promise for gene and cell therapy as well as drug discovery. We next sought to test the knock-in efficiency with cssDNA donor templates. We again compared the knock-in efficiency and cell health of cssDNAs vs dsDNAs as an HDR donor template. In agreement with the previous observations in K562 cells, targeting of the *RAB11A* locus with GFP cssDNA donor templates outperformed dsDNA donor templates with ∼50% knock-in efficiency and without compromising iPS cell health (**Figure 3A and 3B**). Additionally, knock-in efficiency was tested with a range of cssDNA doses. The highest knock-in efficiency (over 60%) was observed with 3 µg of *RAB11A* targeting cssDNA. Similar data was observed for *B2M* targeting cssDNA mediated knock-in, suggesting the versatile application of cssDNA for genome knock-in at different genomic loci (**Figure 3C**). Throughout the process, the engineered iPSC colonies maintained the morphological features that are characteristic of undifferentiated cells (**Figure 3D**). Four cssDNA-engineered iPSC colonies (two with GFP knock-in at the *RAB11A* locus and two with GFP knock-in at the *B2M* locus), when subjected to KaryoStat+ assay, showed no chromosomal aberrations when compared against the reference dataset (**Supplementary Figure 3A**), thus indicating no aneuploidies, submicroscopic aberrations, or mosaic events in cssDNA-engineered iPSC colonies. Furthermore, in a TaqMan hPSC Scorecard Pane assay (**Supplementary Figure 3B**), the same four engineered iPSC colonies had confirmed expression for the nine panel genes associated with self-renewal, while lacking the expression of ectodermal, mesodermal, and endodermal genes, demonstrating that cssDNA engineered iPSC colonies maintain their pluripotency.

**Figure 3.**
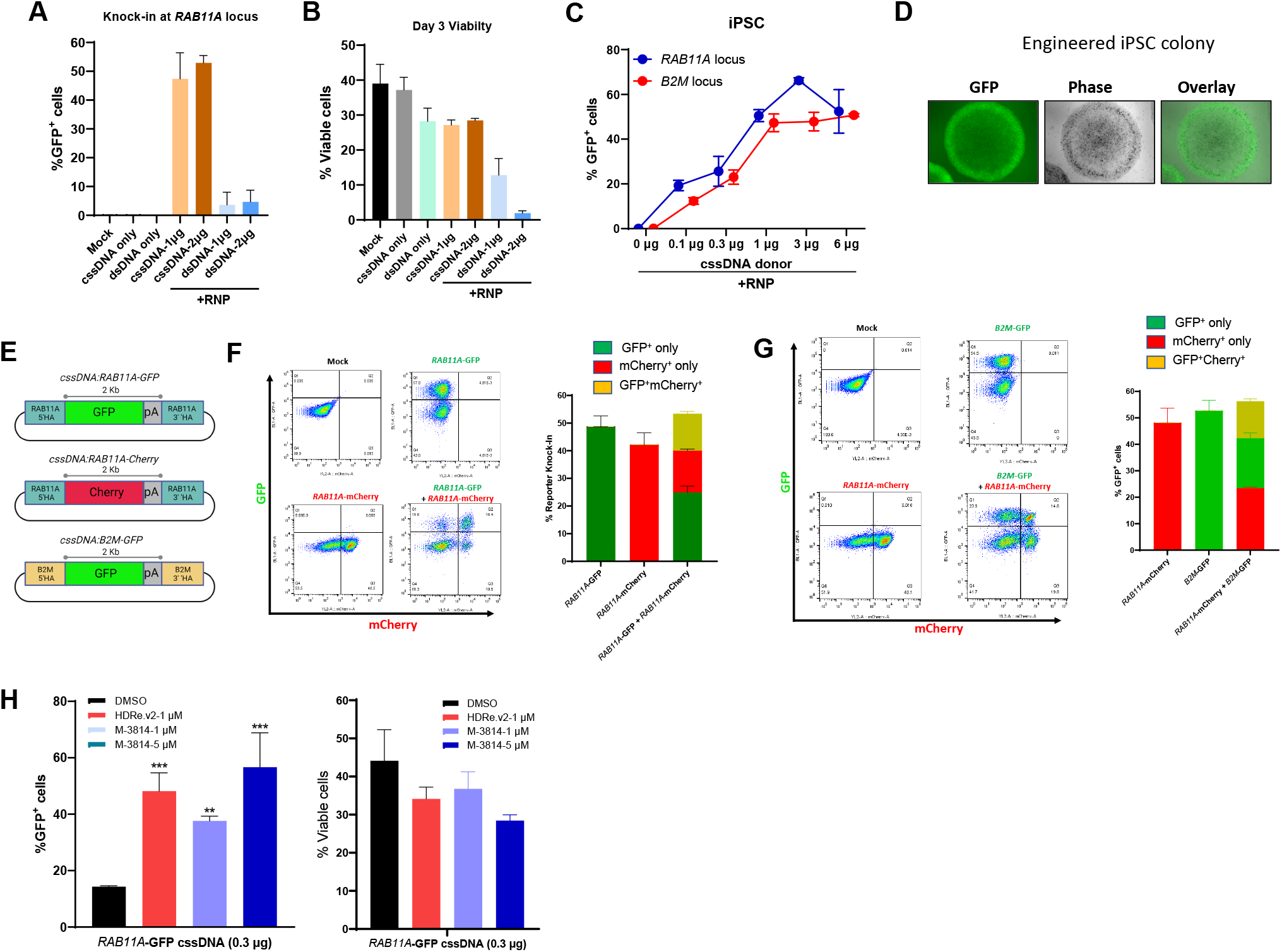
iPSC engineering with cssDNA. **A**. GFP knock-in efficiency (on Day 7) at the *RAB11A* locus in iPSC using cssDNA or dsDNA as donor templates. **B**. Cell viability of cssDNA- and dsDNA-engineered iPSC as measured on Day 3. **C**. Does-dependent GFP reporter knock-in using GFP reporter cssDNA targeting *RAB11A* or *B2M* locus. **D**. GFP fluorescence and phase contrast imaging of representative engineered iPSC colony. **E**. Schematic diagram of cssDNA donor templates for multiplexed reporter gene knock-in at *RAB11A* and *B2M* loci. **F**. Dual knock-in of mCherry and GFP reporter genes at the same *RAB11A* locus. **G**. Dual knock-in of mCherry and GFP reporter genes at *RAB11A* locus and *B2M* locus, respectively. **H**. NHEJ small molecule inhibitors increased genome knock-in efficiency in iPS cells (left) without impacting cell viability (right). iPSCs were treated with NHEJ small molecule at specified concentration for 24 hours. GFP knock-in efficiency was assessed on Day 7 post electroporation by flow cytometry. Cell viability was measured on Day 3 post electroporation. ****, p<0.0001 One-Way ANOVA Bonferroni post hoc test between indicated group.

Bi-allelic genome knock-in at the same locus was previously considered rare, especially for large (multi-kilobase) payload knock-ins. To determine the efficiency of bi-allelic integration, we engineered two donor cssDNA templates with different fluorescence markers (GFP and mCherry) targeting the same *RAB11A* locus site in iPSCs (**Figure 3E**). As expected, when only one reporter cssDNA was delivered with RNP targeting the *RAB11A* locus, 40-50% reporter knock-in were observed. When two reporter cssDNAs were co-delivered with the *RAB11A* RNP in iPSC, ∼25% of cells were GFP only positive and ∼15% of cells were positive for only mCherry, while ∼15% of cells expressed both fluorescence reporters (**Figure 3F**). Expressing both fluorescence reporters indicated bi-allelic integration in the cell. Because the single fluorescence positive cells were expected to contain both mono-allelic and bi-allelic integration populations, a higher percentage (∼50%) of bi-allelic integration at the *RAB11A* locus is expected in the successfully engineered cell population. In addition, the two-fluorescence reporter cssDNAs targeting the *RAB11A* locus or the *B2M* (**Figure 3E**) were designed to determine if simultaneous multiplex knock-in at two different loci can be achieved in iPSC. When both cssDNA donor templates were co-electroporated with two different RNP targeting the *RAB11A* and *B2M* loci, ∼15% of cells expressed both fluorescence reporters (**Figure 3G**). These results demonstrate that cssDNAs are superior HDR donor templates that elicit efficient bi-allelic integration at the same locus and multiplex knock-in at different loci simultaneously.

We next evaluated small-molecule inhibitors that have been reported to enhance knock-in efficiency, including the DNA-dependent protein kinase (DNA-PK) inhibitor M-3814 and “Alt-R HDR enhancer 2” (HDRe.v2), which is described as a nonhomologous end joining (NHEJ) inhibitor (Kath *et al*., 2022; Shy *et al*., 2022; Tatiossian *et al*., 2021). Cells treated for 1 hour after electroporation with either M-3814 or HDRe.v2 (**Figure 3H)** demonstrated increased knock-in efficiency in iPSC cells of up to 200-300% compared to untreated controls. Live cell counts were generally unaffected at the chosen concentrations, except for higher doses of M-3814, which demonstrated a roughly 38% reduction in cell viability at day 4 post-electroporation.

### Efficient Primary B cell and CD34^+^ HSPC Engineering with cssDNA

B cells offer attractive opportunities in gene therapy due to their ability to produce high levels of secreted proteins and persist as long-lived plasma cells (Kleinboehl *et al*., 2022). B cell precision engineering with CRISPR/Cas9 has shown promising efficiency using HDR donor templates delivered by AAV6 (Hung *et al*., 2018; Johnson *et al*., 2018). We attempted to engineer primary B cells using non-viral cssDNAs as HDR donor templates. GFP reporter-tagged dsDNAs or cssDNAs targeting the *RAB11A* locus used in the previous study were co-delivered with an RNP into primary B cells. Knock-in efficiency was determined by flow cytometry for GFP^+^ cells. As shown in **Figure 4A**, B cell engineering with cssDNA HDR donor templates resulted in dose-dependent knock-in, and up to 24% of GFP^+^ cells were achieved with just 3 µg of cssDNA without impacting cell health. However, 3 µg of dsDNA donor templates only resulted in ∼15% of GFP^+^ cells, but cell viability decreased by 80%.

**Figure 4.**
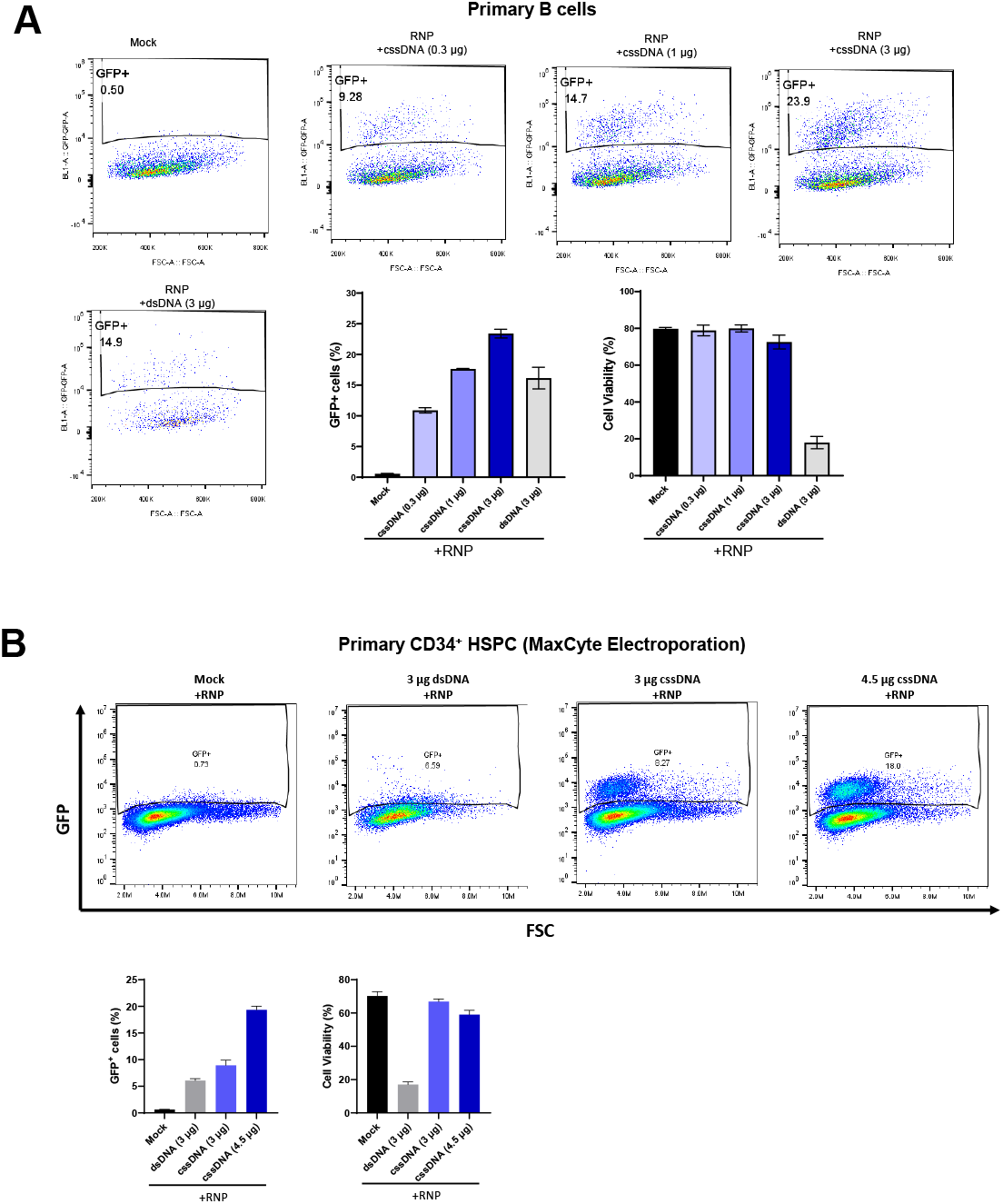
Human primary B cells and CD34^+^ hematopoietic stem and progenitor cells engineering with cssDNA. **A**. GFP reporter knock in efficiency at the *RAB11A* locus in human primary B cells using cssDNA or dsDNA as donor templates. The electroporation was performed in Lonza Amaxa 4D-nucleofector™ system. GFP knock-in efficiency was examined on Day 7 post electroporation by flow cytometry. Cell viability was measured on Day 3 post electroporation. **B**. GFP reporter knock-in efficiency at the *RAB11A* locus in human primary CD34^+^ hematopoietic stem and progenitor cells (HSPC) using cssDNA or dsDNA as donor templates. The electroporation was performed using the MaxCyte ATX system. GFP knock-in efficiency was examined on Day 7 post electroporation by flow cytometry. Cell viability was measured on Day 3 post electroporation.

*Ex vivo* gene therapy based on CD34^+^ hematopoietic stem/progenitor cells (HSPCs) involving lentiviral vectors and NHEJ-based gene-editing has exhibited substantial clinical progress in recent years (*Aiuti et al*., 2013; Cartier *et al*., 2009; Sessa *et al*., 2016). However, studies involving HDR-mediated HSPC engineering, especially with non-viral HDR donor template delivery, have not yet made significant advancements. GFP reporter dsDNA and cssDNA donor templates targeting the *RAB11A* locus used in earlier experiments were co-delivered with an RNP into primary CD34^+^ HSPCs using the MaxCyte ATx system. Knock-in efficiency was determined by flow cytometry for GFP^+^ cells. As shown in **Figure 4B**, up to ∼20% knock-in efficiency was observed with 4.5 µg of cssDNA, and ∼8% of GFP^+^ cells were observed with 3 µg of cssDNA. HSPC viability was generally well maintained for both doses of cssDNA. In contrast, 3 µg of dsDNA donor template only resulted in ∼6% of GFP knock-in, while cell viability was largely compromised.

### cssDNA Efficiently Engineer Primary T Cells without Inducing Innate Cellular Immune Response

Genetically engineered T cell immunotherapies have provided remarkable clinical success in treating B cell acute lymphoblastic leukemia and have the potential to provide therapeutic benefits for numerous other cancers, infectious diseases, and autoimmunity (Ellis *et al*., 2021). Adoptive T cell therapies rely on exogenous gene transfer in primary T cells, resulting in transient or stable transgene expression (Sadelain *et al*., 2017). The five FDA-cleared CAR T cell therapies (Kymriah, Yescarta, Tecartus, Breyanzi, and Abecma) all utilized lentiviral vectors to deliver transgenes into T cells. A variety of non-viral delivery approaches have also been investigated in different stages of preclinical studies. GFP reporter cssDNA HDR donor templates targeting the *RAB11A* locus (*RAB11A*-GFP) or the *TRAC* locus (*TRAC*-GFP) were used to engineer primary T cells. The 2 Kb (excluding the homology arms) *RAB11A*-GFP and 2 Kb or 4 Kb *TRAC*-GFP cssDNA templates were tested in T cell experiments (**Figure 5A**). Primary T cells were co-electroplated with respective RNPs, and knock-in efficiency was determined on Day 7 post electroporation by GFP^+^ cell percentage using flow cytometry. Higher integration efficiency was observed for the 2 Kb payload size compared to the 4 Kb payload. Consistent with findings in iPSCs, DNA-PK inhibitor M-3814 treatment significantly increased knock-in efficiency by 60-150 % (**Figure 5A**).

**Figure 5.**
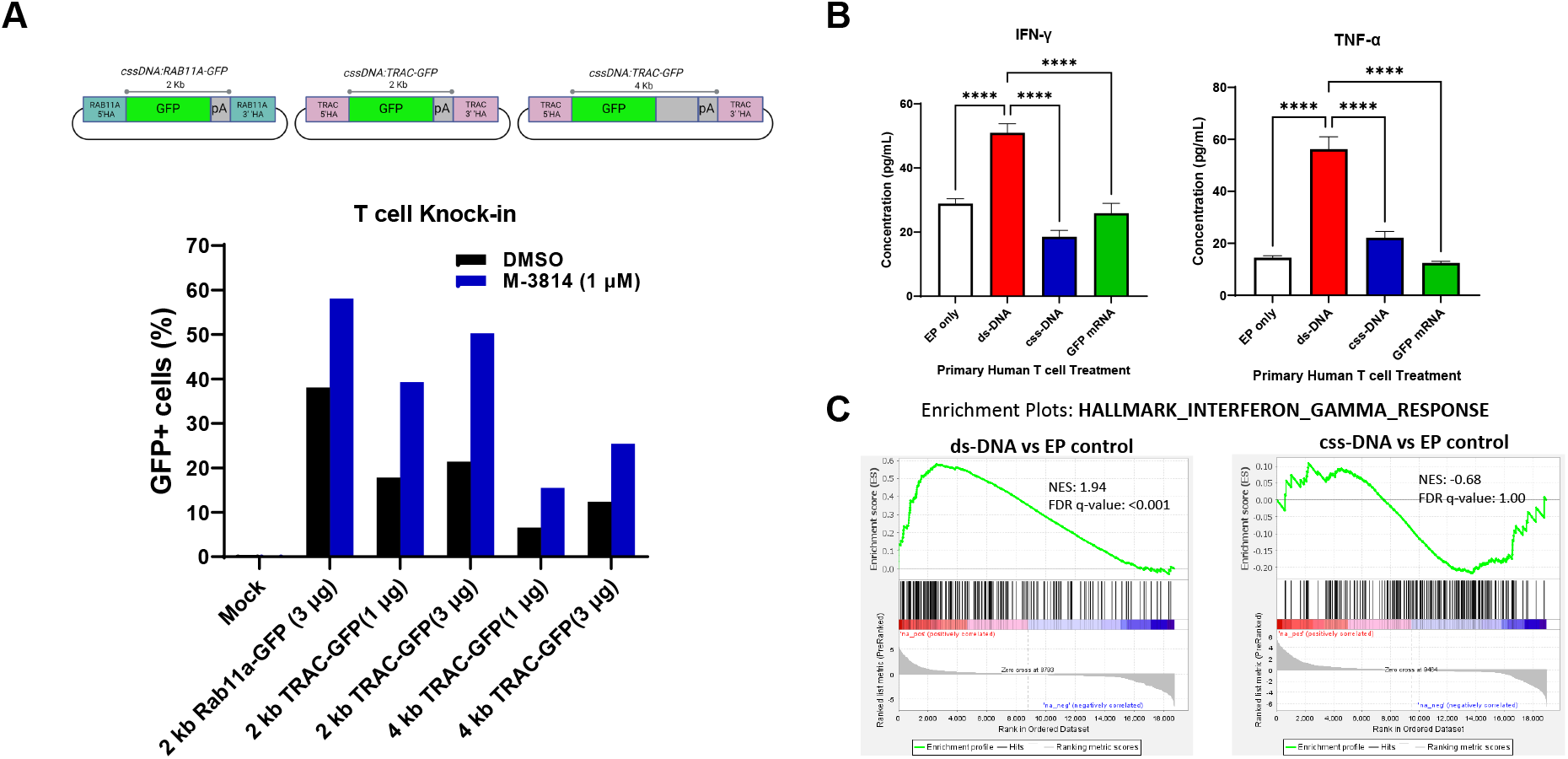
Human primary T cell engineering with cssDNA. **A**. Schematic diagram of the GFP reporter cssDNA targeting *RAB11A* locus and *TRAC* locus (Upper). GFP reporter knock-in efficiency with cssDNA donor templates in human primary T cells. Immediately after electroporation, cells were treated with either DMSO or 1 µM M-3814 for 24 hours. The knock-in efficiency was examined on Day 7 post electroporation. **B**. Secreted IFN-γ and TNF-α cytokines levels in cultured primary T medium after electroporation with buffer, dsDNA, cssDNA or mRNA molecules at 1 µg/million cells. Conditioned media were collected for cytokine analysis using the Ella immunoassay platform with selected panels. **C**. Gene set enrichment analysis (GSEA) analysis of interferon gamma response genes on the differential expression genes (DEG) of RNA samples from dsDNA or cssDNA treated primary T cells. NES: normalized enrichment score; FDR q-value: False Discovery Rate-adjusted p-value. **** p<0.0001 One-Way ANOVA Bonferroni post hoc test between indicated groups.

Traditionally, dsDNAs were used as HDR donor templates for precision genome engineering. However, the usage of dsDNA in gene and cell therapy is largely limited by its cytotoxicity, mainly due to triggering cellular innate immune responses mediated by cytosolic DNA sensing pathways, such as Toll-like receptor 9 (TLR9), cyclic GMP-AMP synthase (cGAS), or stimulator of interferon genes (STING) (Briard *et al*., 2020; Schlee & Hartmann, 2016; Zahid *et al*., 2020). We compared dsDNA- and cssDNA-mediated proinflammatory response in cultured primary T cells. T cells were electroporated with the same amount (1 µg) of dsDNA or cssDNA, GFP mRNA or vehicle (H_2_O). 24 hours post treatment, culture medium was collected for cytokine analysis and cell pellets were collected for transcriptome analysis by RNA sequencing. As shown in **Figure 5B**, interferon-gamma (IFN-γ) and TNF-γ in culture medium were both significantly increased when T cells were treated with dsDNA, but not with cssDNA or mRNA. Consistently, enrichment analysis on the differential expression genes in RNA sequencing analysis demonstrated that dsDNA highly activated intracellular IFN-γ response genes. However, there were no significant changes in IFN-γ response genes when T cells were treated with cssDNA. A panel of innate cellular immune response related genes (OSA1, OSA2, OSA3, OASL, IFIT1, TNF, CCL3, CCL4, etc.) were elevated by dsDNA treatment, but not cssDNA or mRNA (**Supplementary Figure 4**). This demonstrates that unlike dsDNA, cssDNA does not trigger significant cellular proinflammatory response, suggesting it as a much safer HDR donor template.

### cssDNA Donor Template are Compatible with Other Nuclease Editors

Although *Streptococcus pyogenes* Cas9 (SpCas9) is the most widely used CRSIPR variant in genome engineering experiments, its application does have certain limitations, such as a less stringent protospacer adjacent motif (PAM) sequence, which could result in non-specific targeting. The off-target effects might cause detrimental effects in the cells and are especially concerning for clinical applications. Alternative engineered Cas9 variants, such as Cas9 nickases, have been developed by mutating one of the catalytic domains, thus only nicking a single DNA strand at the desired target instead of creating a DSB. Paired nickases targeting opposite complementary strands of DNA, each with a different guide RNA, could be used to create a DSB with high fidelity. The use of paired nickases has been demonstrated to drastically reduce off-target effects while maintaining (or sometimes with higher) Cas9 nuclease efficiency (Cho *et al*., 2014; Gopalappa *et al*., 2018; Ran *et al*., 2013). To evaluate the efficiency of cssDNA-mediated genome knock-in for different variants of Cas9 nucleases, double Cas9 nicking by a pair of sgRNAs was used to introduce targeted DSBs, which has enhanced genome editing specificity (Ran *et al*., 2013). Following the previously reported strategy (Schubert *et al*., 2021), two PAM-out configurations with 41-nt and 57-nt nick distances were designed by D10A SpCas9 nickase (nCas9) (**Figure 6A**) at exon 1 of the *RAB11A* locus. As expected, GFP cssDNA HDR template targeting the *RAB11A* locus did not show significant GFP knock-in when nCas9 RNP with gRNA1, gRNA2 or gRNA alone (**Figure 6B**). Only the two PAM-out configurations with double nCas9 nicking sgRNA1+sgRNA (RNP1+3) or sgRNA2+sgRNA3 (RNP2+3) showed significant GFP knock-in when co-delivered with GFP reporter cssDNA HDR templates in K562 cells (**Figure 6B**). RNP2+3 pair showed significantly higher efficiency than RNP1+3, although lower efficiency compared to wild type spCas9 (wtCas9) with RNP, which induced blunt end DSBs. As a control, RNP1+2, which both nicked the same targeting strand, did not induce any knock-in. This demonstrates that cssDNA donor templates are compatible with different Cas9 variants to provide a versatile toolbox for efficient and safe genome knock-in.

**Figure 6.**
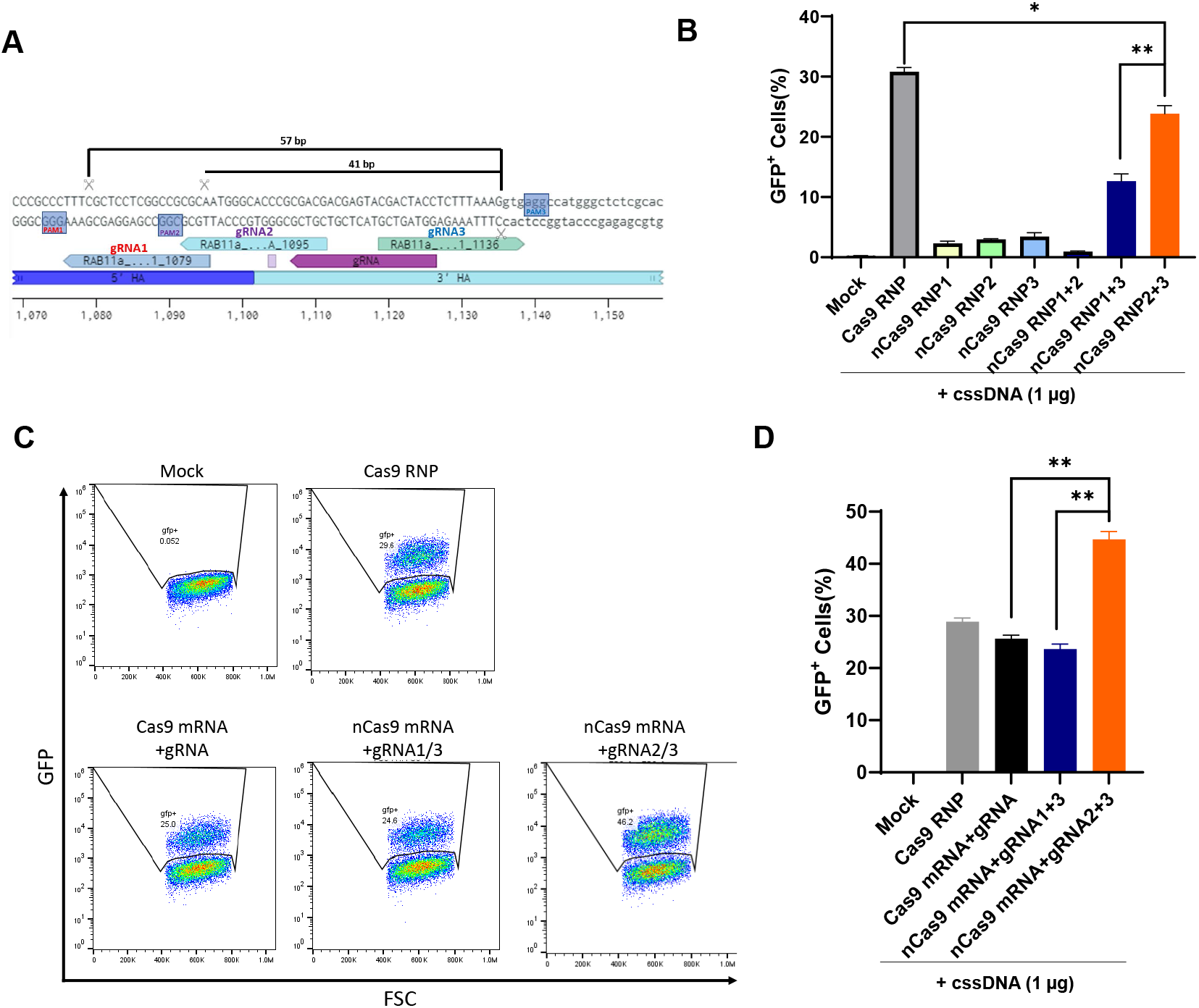
K562 genome engineering with paired spCas9 nickases and cssDNA. **A**. Schematic diagram of sgRNA design for paired Cas9 nicking at exon 1 of the *RAB11A* locus. Two PAM-out configurations were designed for the double Cas9 nicking (gRNA2+3 and gRNA2+3) with 57 bp and 41 bp apart between the nicking sites. **B**. GFP knock-in efficiency at *RAB11A* locus using cssDNA donor template with paired Cas9 nicking design. Knock-in efficiency was examined on Day 7 post electroporation by flow cytometry. **C**. Representative GFP flow cytometry graphs of engineered K562 cells with Cas9 or D10ACas9 (nCas9) mRNA delivery. **D**. Quantification of the GFP knock-in efficiency when Cas9 or nCas9 was delivered by mRNA. ** p<0.01, One-Way ANOVA Bonferroni post hoc test between indicated groups.

We further tested the efficiency when delivering Cas9 by mRNA rather than protein (RNP). Similar knock-in efficiency was observed with Cas9 RNP and Cas9 mRNA (Figure 6C and 6D). When gRNA1+3 or gRNA2+3 was co-delivered with nCas9 mRNA, the knock-in efficiency was significantly higher than when nCas9 was delivered by protein (RNP) (**Figure 6C and 6D**). nCas9 mRNA gRNA2+3 was more efficient than Cas9 mRNA gRNA for cssDNA-mediated knock-in. These results indicate that GATALYST gene writing using cssDNA HDR donor template is agonistic to various nucleases delivered in different forms (protein or mRNA). Furthermore, mRNA delivery could result in more durable Cas9 expression, and is expected to have a higher nuclease activity to induce two-nicking DSBs compared to protein delivery. Our data aligns with this, as cssDNA HDR donor-mediated knock-in is significantly higher than RNP delivery, further supporting cssDNA as a superior editing tool.

### Efficient CAR-T Engineering with Durable Cancer Cell Lysis Function using cssDNA Donor Template

Finally, we sought to engineer T cells with CARs using non-viral cssDNA donor templates to demonstrate the therapeutic potential of cssDNAs (**Figure 7A**). We engineered bi-specific CARs targeting CD19 and CD22 (Qin *et al*., 2018; Spiegel *et al*., 2021), which offered great potential to treat CD19 CAR resistance relapsed or chemotherapy-refractory (relapsed/refractory) B cell malignancy resistance, into the endogenous *TRAC* locus. Targeting a CAR to the *TRAC* locus greatly enhanced the potency of CAR-T cells in preclinical studies and delayed effector T-cell differentiation and exhaustion (Eyquem *et al*., 2017). Anti-CD19xCD22-CAR cssDNA targeting the *TRAC* locus was co-electroporated with spCas9 protein and *TRAC* sgRNA. Day 7 post electroporation, CAR expression was determined by Protein L binding as previously described (Zheng *et al*., 2012). Approximately 33% knock-in efficiency (percentage of CAR+ cells) was achieved with 2 µg of cssDNA donor templates (**Figure 7B**). When treated with DNA-dependent protein kinase inhibitor M-3814, knock-in efficiency further increased to 56.3% (**Figure 7B**). Similar data was obtained with CD19 CAR specific detection reagent for CAR detection in flow cytometry (data not shown). Protein L was used for all the following CAR detection experiments. To benchmark against the engineering with donor template delivered by rAAV in previous studies, equivalent *TRAC* locus cssDNA and AAV6 vectors encoding an anti-CD19xCD22-CAR were made, and direct comparisons demonstrated efficient knock-in with both approaches. Although, consistently higher CAR expression efficiency was observed with AAV6 vector templates (75-80%) than with cssDNA templates (40-50%) (**Figure 7C**). Both cssDNA and AAV6 engineered T cells expanded 25-30-fold during 7 days of *ex vivo* culture, which was only slightly lower than un-engineered mock T cells (**Figure 7D**). On Day 7 post engineering, the cytotoxicity of engineered CAR-T cells was examined. *In vitro* assays demonstrated efficient CAR-T cell killing of CD19^+^ and CD22^+^ NALM6 cells (**Supplementary Figure 5**), in contrast to the mock engineered T cells from the same donor (**Figure 7E**). Interestingly, although at lower knock-in efficiency, the target cell killing activity of cssDNA engineered CAR-T was significantly higher than by the AAV6 engineered CAR-T cells at 24-hour time point, suggesting higher potency of cssDNA CAR-T cells than AAV6 CAR-T cells. Furthermore, when *in vitro* cell killing effects were monitored for extended time (96 hours), target cells re-grew significantly with un-engineered T cells as well as with AAV6 engineered CAR-T cells at most conditions except 6:1 effector:target ratio in the killing assay (**Figure 7F**). However, target cell re-growth was largely prohibited by cssDNA CAR-T cells at all the tested E:T ratios. This demonstrates that despite lower CAR knock-in efficiency, ccsDNA engineered CAR-T cells present with higher potency and higher durability against CD19^+^CD22^+^ target B cell leukemia cells.

**Figure 7.**
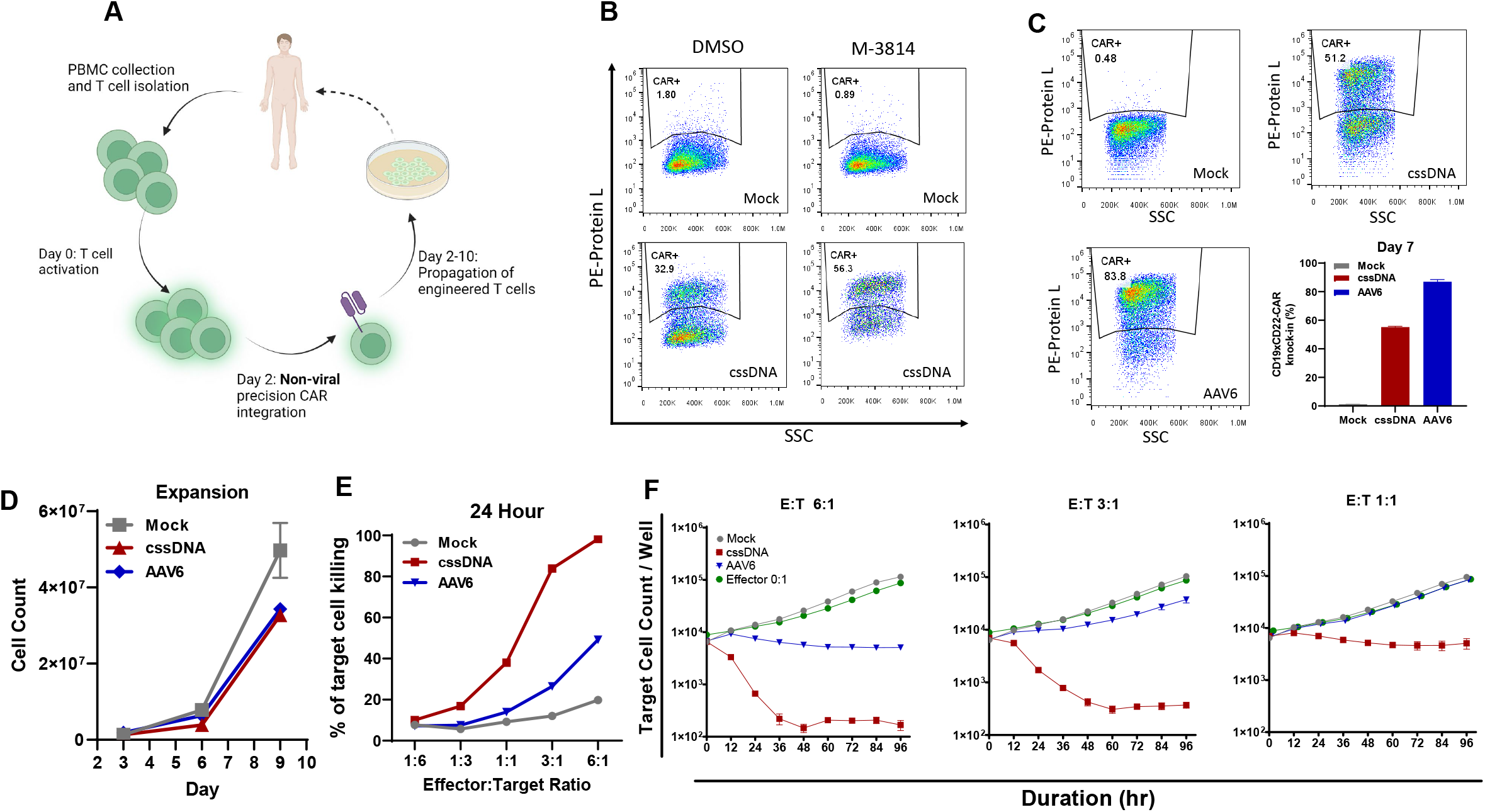
CAR-T cell engineering with cssDNA. **A**. Schematic diagram of a nonviral CAR-T cell engineering process. Pan T cells were isolated from peripheral blood and activated on day 0 with anti-CD3/anti-CD28 TransAct. Cells were electroporated using the Lonza nucleofector on day 2 with Cas9 RNPs + cssDNA HDR donor templates and then expanded for a total of 7–10 days. **B**. Representative Day 7 flow cytometry graphs showing CAR knock-in for Mock (un-engineered) and cssDNA-engineered T cells with DMSO or 1 µM M-3814 treatments. **C**. Representative Day 7 flow plots and quantifications of CAR knock-in for mock (un-engineered), cssDNA- or AAV6-engineered T cells. Cells were treated with 1 µM M-3814 for 1 day immediately after electroporation. **D**. Expansion ability of engineered CAR-T cells. **E**. *In vitro* killing of NALM6 acute lymphoblastic leukemia cell line with cssDNA-, AAV6-engineered CAR-T cells in comparison to unmodified T cells from the same blood donor after 24 hours of co-culture. The *in vitro* killing measured by live cell imaging using the IncuCyte® live cell imaging system. **F**. The growth curve of target NALM6 cells when co-cultured with un-engineered, cssDNA- or AAV6-engineered CAR-T cells at various effector to target (E:T) ratios.

## Discussion

Single-stranded DNA (ssDNA) exists in nature, which only commonly occurs in viruses or in a transient form during dsDNA replication process. Synthetic biology is an emerging field of genome engineering with specific purposes in medicine, manufacturing, and agriculture. In the past several years, ssDNA has attracted attention across the genetic engineering field, mainly due to its low cellular toxicity and high insertion efficiency in CRISPR/Cas-based genomic editing (Codner *et al*., 2018; Dokshin *et al*., 2018; Du *et al*., 2018; Iyer *et al*., 2022; Lanza *et al*., 2018; Li, 2019; Miura *et al*., 2018; Roth *et al*., 2018; Xiao *et al*., 2018). Although lssDNA can be chemically synthesized or produced *in vitro* by enzymatic processes, these methods can be costly, lack scalability, and currently cannot create long molecules (over 5 Kb). In addition, the produced linear strands are prone to exonuclease degradation (Veneziano *et al*., 2018). The manufacturing of high purity cssDNAs up to 20 Kb in length from engineered phagemids provides a low-cost alternative. cssDNAs are more resistant to exonuclease activity, which may prove important for therapeutic applications.

Thus, the new technological developments in synthetic cssDNA production can be made bio-orthogonal and scalable, enabling future applications to novel therapeutics. Our current cssDNA purification process from engineered phagemid is research grade at small scale with relatively low yield. However, its production was proven to be amendable to a scalable bath fermenter (Shepherd *et al*., 2019).

The yield of cssDNA varied from 0.2 mg to 4 mg per L of culture during our development process, and we reasoned that the low yield of certain cssDNA templates might be attributed to specific DNA molecule sequence because ssDNA molecules do not pair up to form helices and are prone to form secondary structures or topologically linked circular molecules (Liang e*t al*., 2006). These secondary structures could impact the cssDNA packaging into phage particles. Nevertheless, those cssDNAs with intrinsically low yield can be successfully manufactured with high purity. Higher yield that meets future clinical application is expected with the use of a scalable fermenter culture system.

For targeted genome editing, most previous non-viral approaches utilized oligonucleotides, plasmids, or linear dsDNAs as the donor templates to achieve targeted DNA insertion at a locus of interest (Salsman & Dellaire, 2017). lssDNAs were demonstrated as superior donor templates because of the reduced cellular toxicity and increased HDR efficiency compared to dsDNA (Bai *et al*., 2020; Li, 2019; Miura *et al*., 2018; Quadros *et al*., 2017; Shy *et al*., 2022). Moreover, with the use of ssDNA, illegitimate random integration is expected to be less frequent as opposed to the use of dsDNA templates (Won & Dawid, 2017; Wurtele *et al*., 2003; Zorin *et al*., 2005). The lssDNA donors used in these seminal studies were exclusively linear molecules produced with *in vitro* methods by manipulation of DNA with polymerase chain reaction coupled with enzymatic degradation of one of the two DNA strands (Li, 2019; Shy *et al*., 2022). However, the enzymatic production of lssDNA templates is inherently inefficient and only up to 5 Kb ssDNA is commercially offered due to the accumulation of mutations during PCR, expensive reagent cost, and low scalability. These limitations of lssDNA are therefore prohibitive for clinical application.

Accumulating evidence suggests that genome integration with the existing CRISPR/Cas9-mediated HDR technology in engineered cells is often mono-allelic (He *et al*., 2016; Wang *et al*., 2016; Ye *et al*., 2014). However, for applications in monogenic disease mutation correction, gene therapy, transgenic model animal generation, and agricultural manufacturing, biallelic genome modification is highly desirable. cssDNA is a superior HDR donor template, not only due to its low toxicity and high knock-in efficiency, but also for its high level of biallelic integration, allowing its application to many versatile applications in basic research, gene and cell therapy, agriculture, and climate change.

Overall, phagemid-derived cssDNAs were demonstrated to be superior HDR donor templates offering highly efficient genome knock-in with low cellular toxicity. cssDNA production is bio-orthogonal and scalable, with the capability to generate cssDNA lengths well beyond the packaging capacity of current viral vectors. cssDNAs were shown to be well suited for use with multiple nucleases, in multiple cell types at multiple genetic loci. In immune cell therapy, ∼ 50% targeted CAR knock in on the *TRAC* locus has been proved using cssDNAs as HDR donor templates in primary T cells. cssDNA engineered CAR-T cells showed higher potency and higher durability to target tumor cells when compared to CAR-T cells engineered with donor templates delivered by AAV6. These advantages make cssDNA an attractive alternative for HDR donor templates for efficient insertion of large DNA payloads in a variety of disease-relevant cell types and can be leveraged for basic research and gene and cell therapies.

## Materials and Methods

### Generation of HDR template circular single-stranded DNA from M13 phage

Donor template sequences (transgene sequence flanked with 5’ and 3’ homology arms at 300-500 nt in length) are constructed as dsDNA and cloned into phagemid vector. An XL1-Blue *E. coli* strain was co-transformed with the M13 helper plasmid and phagemid containing double-stranded donor template and selected on agar plates with kanamycin (50 µg/mL) and carbenicillin (100 µg/mL). A single colony was selected and grown for ∼24 hours (37 °C, 225 rpm) in 250 mL 2xYT media (1.6% tryptone, 1% yeast extract, 0.25% NaCl) to reach OD_600_ between 2.5-3.0. The bacteria were pelleted by centrifugation and the phage particles in the supernatant were precipitated with PEG-8000. The precipitated phage particles were then pelleted by centrifugation, washed, and lysed in 20 mM MOPS., 1M Guanidine-HCl and 2% Triton X-100. The cssDNA released from the phage were then extracted with NucleoBond Xtra Midi EF kit (Macherey-Nagel) following the manufacturer’ s instructions. The concentration of cssDNA was determined by Nanodrop for ssDNA and the yields were 10 µg per ml of liquid culture. Ratios of Absorbance (A_260_ nm/A_280_ nm and 260nm/230nm) reflect consistent purity (1.8 and > 2, respectively) from serial preps. Recombinant cssDNA is verified by Sanger DNA sequencing using custom-designed staggered sequencing primers for complete coverage.

### Cell culture

K562 (ATCC) and NALM6 (ATCC) cells were maintained in RPMI-1640 media with 10% FBS and 1% penicillin and streptomycin. HEK293T (ATCC) cells were cultured in Dulbecco’ s modified Eagle’ s medium supplemented with 10% fetal bovine serum (FBS) and 1% penicillin and streptomycin (Gibco). iPSCs (ThermoFisher Scientific) were cultured in complete StemFlex (ThermoFisher Scientific) media in vitronectin-coated flasks. iPSC colonies were checked regularly and passaged using ReLeSR (StemCell Technologies) every 3-4 days of culture. iPSCs were ready for electroporation after 2-3 passages. Human primary CD34^+^ HSPCs were isolated from fresh Leukopak using a CD34 MicroBead Kit in MultiMCAS Cell24 Separator Plus (Miltenyi). The cryopreserved HSPCs were thawed, washed, and cultured in StemSpan SFEM II medium supplemented with StemSpan™ CD34+ Expansion Supplement and UM729 (StemCell Technologies). HSPCs were ready for electroporation after 4-5 days in culture. Human primary B cells were isolated using a StraightFrom Leukopak CD19 MicroBead Kit in MultiMCAS Cell24 Separator Plus (Miltenyi). Cryopreserved B cells were thawed, washed, and cultured and activated in ImmunoCult™-XF B Cell Base Medium supplemented with ImmunoCult™-ACF Human B Cell Expansion Supplement (StemCell) for 4 days before electroporation. Human primary T cells were isolated using a StraightFrom Leukopak CD4/CD8 MB Kit in MultiMCAS Cell24 Separator Plus. T cells were cultured and expanded in TexMACS Medium (Miltenyi) supplemented with 200 IU/mL Human IL-2 IS (Miltenyi). T cells were activated for 2 days with T Cell TransAct (Miltenyi) before electroporation.

All cells were maintained in a humidified incubator at 37 °C and 5 % CO_2_, unless otherwise specified. Cell count viability was determined using a Via2-Cassette in NucleoCounter® NC-202 (ChemoMetec) on specified days after engineering.

### Electroporation

All HEK293T, K562, iPS, B and T cell electroporations were performed using the Amaxa™ 96-well Shuttle™ with the 4D Nucleofector (Lonza). HSPC electroporations were performed in MaxCyte ATx. 25 picomole of sNLS-SpCas9-sNLS Nuclease (Aldevron) or D10A Cas9 nickase (nCas9) (ThermoFisher) along with 50 picomol of sgRNA (for SpyCas9, synthesized at Integrated DNA Technologies) were used per reaction. Cas9 nucleases and sgRNAs were precomplexed in supplemented Nucleofector® Solution for 20 min at room temperature and the RNP solution was increased to a final volume of 2.5 µL (10X) per electroporation reaction. For mRNA delivery nucleases, 1 µg of Cas9/nCas9mRNA was co-electroplated with 50 picomol of sgRNA. For electroporating K562 cells, an SF Cell Line 4D-Nucleofector™ Kit and 250,000-500,000 cells per reaction were used. For electroporating HEK293T cells, an SF Cell Line 4D-Nucleofector™ Kit and 200,000-300,000 cells per reaction were used. For electroporating iPSC cells, 100,000 cells per reaction were used with a P3 Primary Cell 4D-Nucleofector™ Kit. For electroporating primary B cells, 1,000,000 cells per reaction were used with a P3 Primary Cell 4D-Nucleofector™ Kit.

For electroporating primary T cells, 2,000,000 cells per reaction were used with a P3 Primary Cell 4D-Nucleofector™ Kit. For electroporating primary HSPC cells, 1,250,000 cells per reaction were used with a 50 µL reaction cuvette in MaxCyte ATx. The indicated amount of HDR donor template (cssDNA or dsDNA) were used in co-electroporation with RNP or Cas9/nCas9 mRNA and sgRNA. After electroporation, the cells were placed in a humidified 32°C incubator with 5 % CO2 for 12-24 hours, followed by transferring to 37°C incubator for 3-10 additional days.

### Quantification of dsDNA or cssDNA by qPCR from electroporated K562 cells

Following from 1-10 days after electroporation, K562 cells were pelleted, and total DNA was isolated using a PureLink Genomic DNA Mini Kit (ThermoFisher Scientific). 20 ng of DNA was used for quantification of cssDNA or dsDNA using a TaqMan Multiplex Master Mix following the manufacturer’ s recommendations. Amplification was detected in a QuantStudio 12K Flex Real-Time PCR System instrument from ThermoFisher Scientific. cssDNA and dsDNA specific TaqMan probes purchased from IDT are shown in **Supplementary Table 1**. The relative remaining DNAs were normalized using 2^ΔCt^ method.

### Flow cytometry analysis

All flow cytometry was performed on an Attune NxT flow cytometer with a 96-well autosampler (ThermoFisher Scientific). Unless otherwise indicated, cells were collected 4-7 days post electroporation, resuspended in fluorescence-activated cell sorting (FACS) buffer (2% FBS in PBS) and stained with 7-AAD (BioLegend), and the indicated cell-surface marker. To obtain comparable live cell counts between conditions, events were recorded from an equivalent fixed volume for all samples. Data analysis was performed using FlowJo_v10.8.0_CL software with exclusion of subcellular debris, singlet gating and live:dead stain. Analyzed graphs were produced with Prism 9 (GraphPad).

### GFP-NALM6 stable cell line generation

NALM6-GFP stable cell line were generated by knock-in EF1a-GFP in the *AAVS1* locus using cssDNAs as donor templates. NALM6 cells were electroporated using an Amaxa™ 96-well Shuttle™ in 4D Nucleofector with 25 pmol of Cas9 and 50 pmol of gRNA targeting the *AAVS1* locus, and 2 µg of cssDNA containing EF1α driven GFP sequence flanked with *AAVS1* 5’ and 3’ homology arms.

Engineered cells were cultured for 3-weeks and flow sorted to dispense single GFP-positive cells into individual wells of 96-well plates in a BD FACSAria™ III (BD Biosciences). NALM6-GFP stable cells from a single colony were expanded. NALM6-GFP single clone stable cells were used in *in vitro* cell cytotoxicity assays.

### CAR-T cell engineering

48 hours after initiation and activation, T cells were electroporated using an Amaxa™ 96-well Shuttle™ in 4D Nucleofector. 2×10^6^ cells were mixed with 25 pmol of Cas9 and 50 pmol of sgRNA (RNP) into each well. For cssDNA engineered cells, 2 µg of cssDNA encoding bi-specific CD19×CD22 CAR was electroporated with an RNP targeting the *TRAC* locus. For AAV6-medaited engineering, recombinant AAV6 donor vector (manufactured by PackGene) was added to the culture immediately after electroporation at 2 × 10^4^ multiplicity of infection. Following electroporation, cells were diluted into culture medium in the presence of T Cell TransAct with DMSO or 1 µM M-3814, and incubated at 32 °C, 5% CO_2_ for 24 hours. Cells were then washed and subsequently transferred into a G-Rex 24 Multi-Well Cell Culture Plate (Wilson Wolf) in standard culture conditions at 37 °C, 5% CO_2_ in IL-2 (200 IU/mL) supplemented TexMACS medium and replenished every 3-4 days. CAR expression was determined using PE labelled Protein L (ACROBiosystems), or Biotin conjugated CD19 antigen fused recombinant human IgG1 Fc (Miltenyi) in PBS with 2 % BSA before flow analysis.

### *In vitro* cell killing

The cytotoxicity of T cells transduced with a bi-specific CD19xCD22 CAR was determined by IncuCyte Cytotoxicity Assay with Annexin V Red Reagent for apoptosis (Sartorius) with NALM-6 cells that stably express GFP serving as target cells. On Day 7-9 post electroporation, the CAR-T effector (E) and tumor target (T) cells were co-cultured in triplicates at the indicated E/T ratio using flat bottom poly-L-ornithine (Sigma) coated 96-well plates with 1 × 10^4^ target cells in a total volume of 200 µL per well in NALM-6 medium. The number of effector CAR T cells was calculated based on the percentage of CAR-positive cells. Target cells alone were plated at the same cell density to determine cell proliferation. Plates were incubated at 37°C with 5 % CO_2_ for up to 4 days. Four images were recorded per well every 2 h and analyzed using IncuCyte cell by cell analysis software module (Sartorius). The killing potency of the CAR-T cells was assessed by comparing the percentage of Annexin V positive target cells over time relative to the total number of target cells.

## Acknowledgments

We thank Dr. Taegon Cha for designing and cloning of initial phagemid constructs and the guidance and suggestions on cssDNA production. We also thank Dr. Matthew C. Lorence for comments and suggestions on the manuscript.

