## Supplementary Figure 1 for "Circular single-stranded DNA is a superior homology-directed repair donor template for efficient genome engineering"

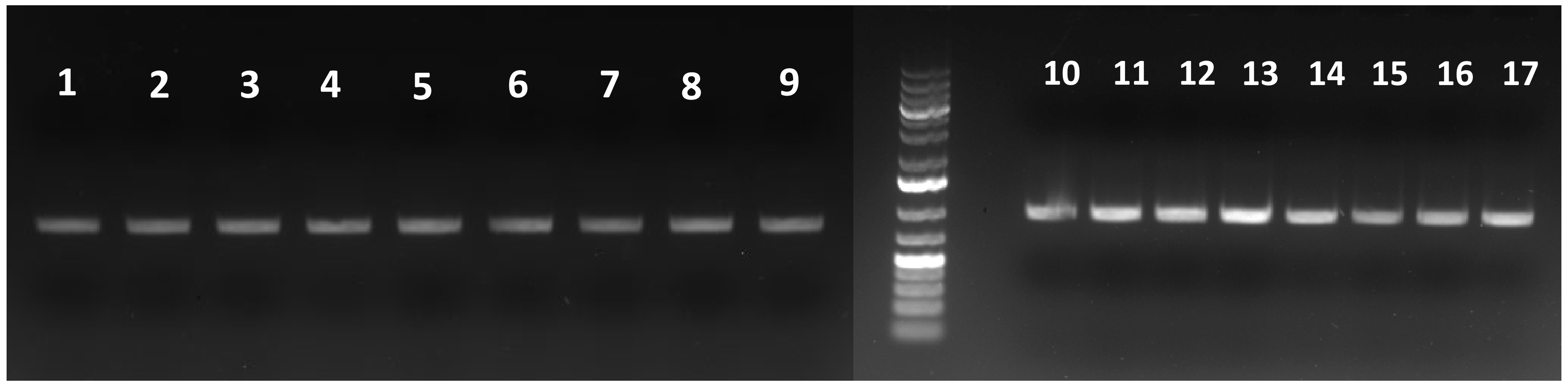


1. Room temperature-1d
2. Room temperature-3d
3. Room temperature-7d
4. Room temperature-10d
5. Room temperature-14d
6. 4°C-1d
7. 4°C-3d
8. 4°C-7d
9. 4°C-10d
10. 4°C-14d
11. Freeze-thaw 3 times
12. Freeze-thaw 9 times
13. Freeze-thaw 15 times
14. Freeze-thaw 21 times
15. Freeze-thaw 30 times
16. Freeze-thaw 42 times
17. -20°C storage

**Supplementary Figure 1. Stability of purified cssDNA under various storage conditions.** cssDNA was aliquoted and stored at room temperature, 4 °C for up to 14 days (lanes 1-10) or was subjected to freeze-thaw cycles for up to 42 times over 14 days (lanes 11-16). An aliquot of cssDNA was stored at -20 °C for 14 days for comparison (lane 17). 100 ng of cssDNA was loaded into DNA agarose gel for imaging.
