## Supplementary Figure 2 for "Circular single-stranded DNA is a superior homology-directed repair donor template for efficient genome engineering"

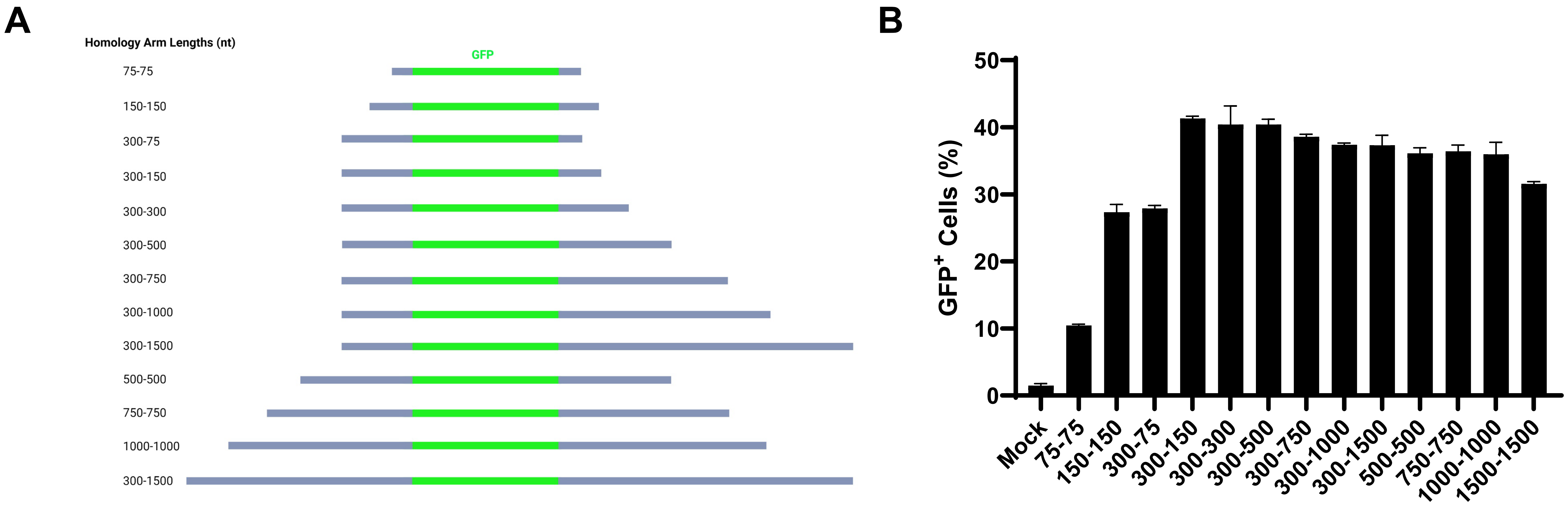


**Supplementary Figure 2. Knock-in efficiency with cssDNAs harboring different lengths of homology arms.** Schematic diagram of GFP cssDNA targeting *RAB11A* locus flanked with different 5' and 3' homology arms. GFP knock-in efficiency with different cssDNA with various homology arm lengths. 500,000 K562 cells were co-electroporated with RAB11A RNP (spCas9 and sgRNA) and 1.8 pmol of cssDNA. Knock-in efficiency was determined by flow cytometry on Day 4 post electroporation.
