## Supplementary Figure 3 for "Circular single-stranded DNA is a superior homology-directed repair donor template for efficient genome engineering"

**A**

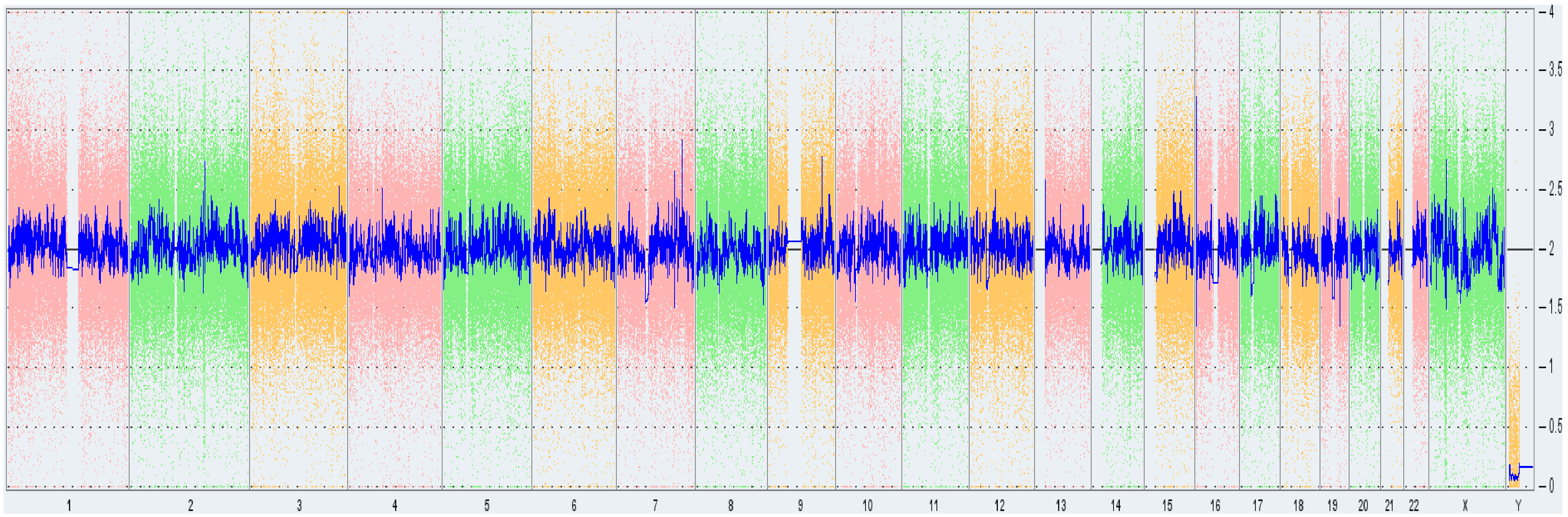


**B**

| Sample Name | Self-renewal | Ectoderm | Mesoderm | Endoderm |
|---|---|---|---|---|
| SC 1449 | 0.41 | -0.53 | -0.39 | -1.01 |
| SC 1450 | 0.09 | -0.55 | -0.85 | -1.33 |
| SC 1451 | -0.10 | -0.64 | -0.50 | -1.15 |
| SC 1452 | 0.08 | -0.57 | -0.74 | -1.51 |
| SC 1453 | -0.44 | -0.61 | -0.03 | -1.22 |

Gene expression relative to the reference standard

| Upregulated | | | | | | Downregulated |
|---|---|---|---|---|---|---|
| x > 1.5 | 1.0 < x ≤ 1.5 | 0.5 < x ≤ 1.0 | -0.5 ≤ x ≤ 0.5 | -1.0 ≤ x < -0.5 | -1.5 ≤ x < -1.0 | x < -1.5 |

**Supplementary Figure 3. KaryoStat+ and Scorecard analysis of cssDNA-enginerred iPSC colones.**
A. Whole genome view of KaryoStat+ data from representative enginerred-iPSC clone. The whole genome view displays all somatic and sex chromosomes in one frame with high level copy number. The smooth signal plot (right y-axis) is the smoothing of the log2 ratios which depict the signal intensities of probes on the microarray. A value of 2 represents a normal copy number state (CN = 2). A value of 3 represents chromosomal gain (CN = 3). A value of 1 represents a chromosomal loss (CN = 1). The pink, green and yellow colors indicate the raw signal for each individual chromosome probe, while the blue signal represents the normalized probe signal which is used to identify copy number and aberrations (if any).*
B. Scorecard Values of un-engineered control iPSC clone (SC 1449) and four cssDNA-engineered clones. Algorithm scores for the samples show upregulation or downregulation of the endoderm, mesoderm, ectoderm or pluripotent (self-renewal) markers relative to the reference set of nine undifferentiated pluripotent stem cell lines.
