## Supplementary Figure 4 for "Circular single-stranded DNA is a superior homology-directed repair donor template for efficient genome engineering"

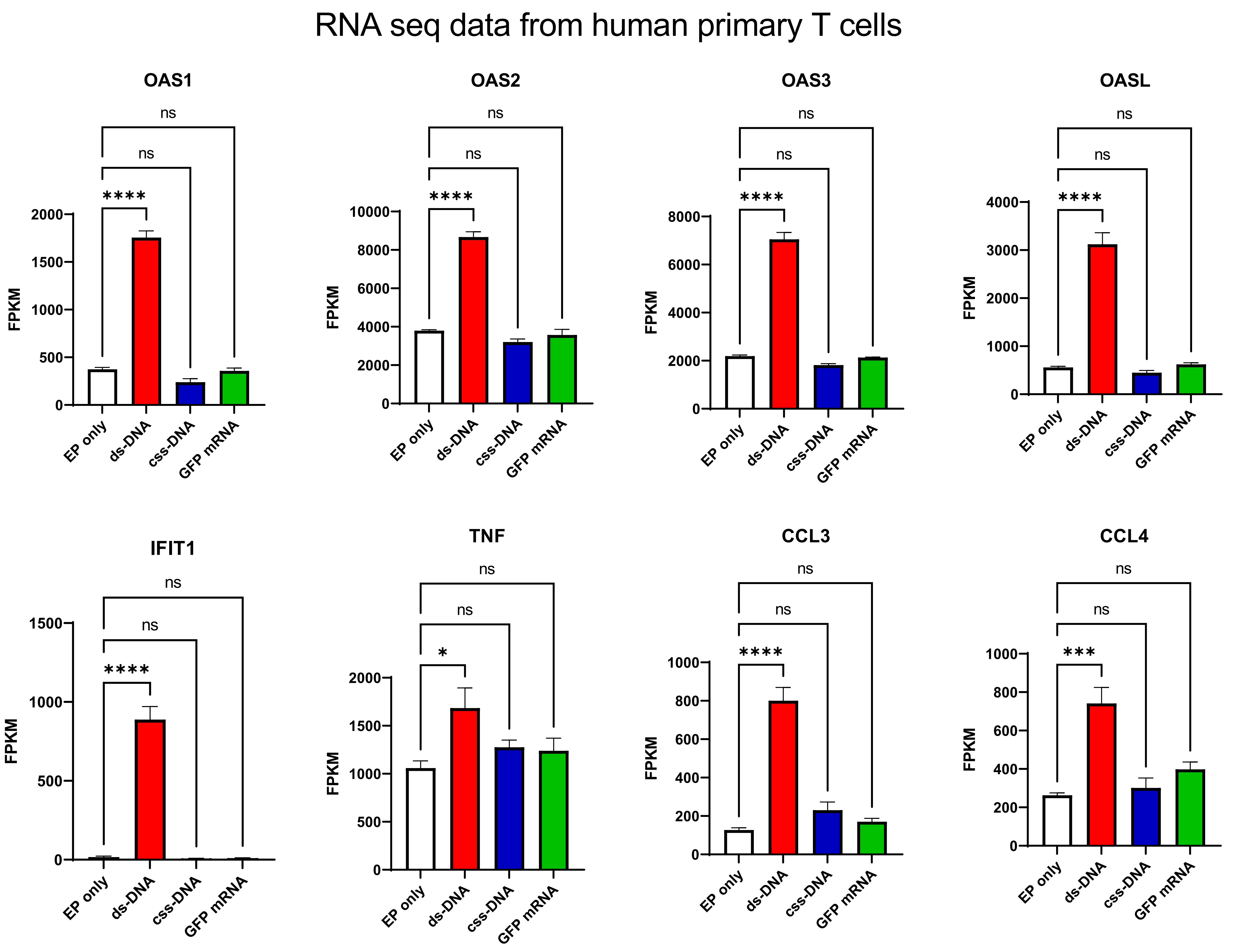


**Supplementary Figure 4. Innate cellular immune response related gene expression level in human primary T cells.** Cultured human primary T cells were electroporated with buffer, dsDNA, cssDNA or mRNA. Cells were collected 24 hours post electroporation for RNA extraction and sequencing. Innate cellular immune response related gene panel were plotted for control (EP only), dsDNA, cssDNA or mRNA treated groups. FPKM, fragments per kilo base of transcript per million mapped fragments. ns, non-significant difference. *, $p<0.05$, ***, $p<0.001$, ****, $p<0.0001$ One-Way ANOVA Bonferroni Post Hoc Test between indicated groups.
