## Supplementary Figure 5 for "Circular single-stranded DNA is a superior homology-directed repair donor template for efficient genome engineering"

**A**

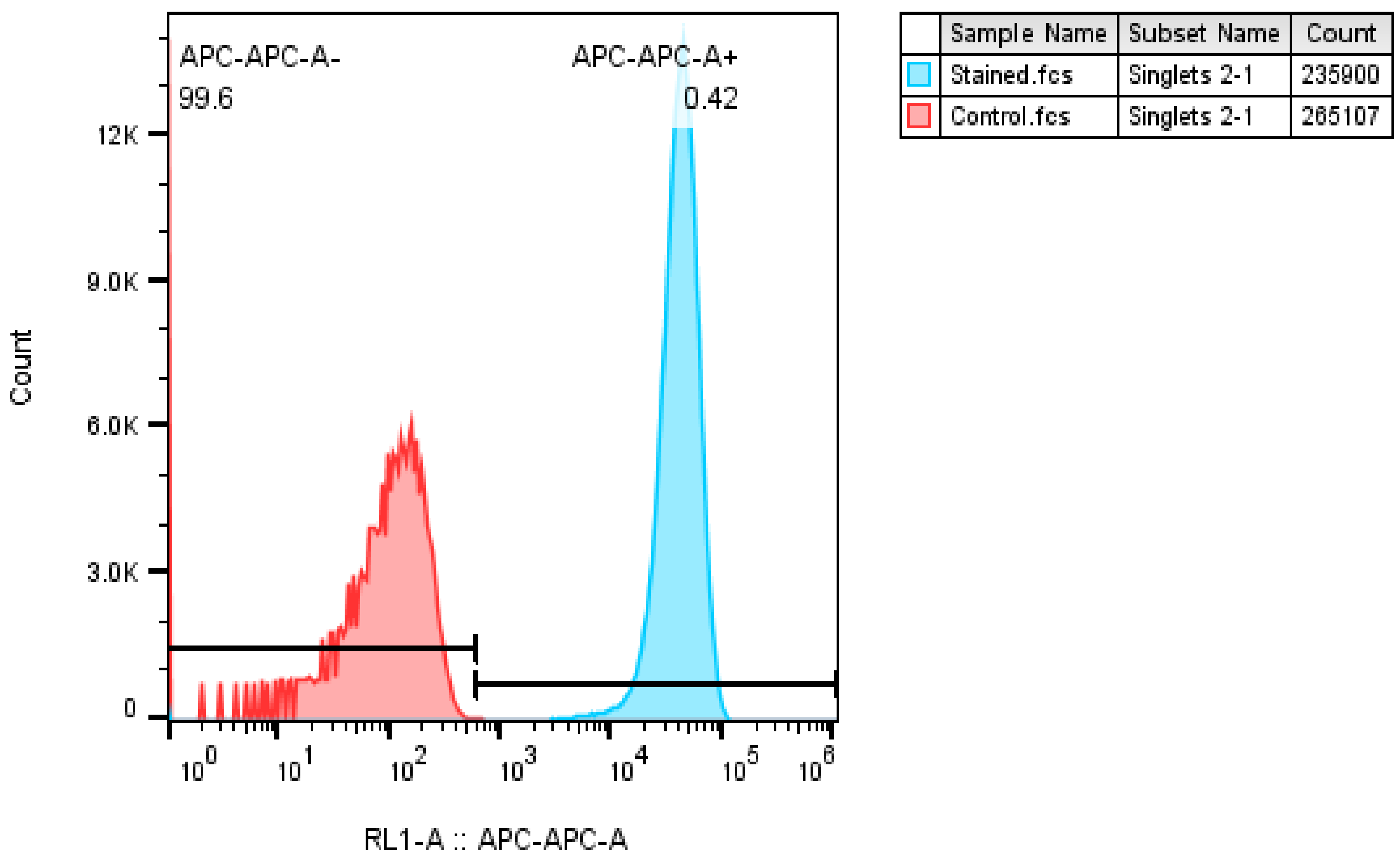


CD19

**B**

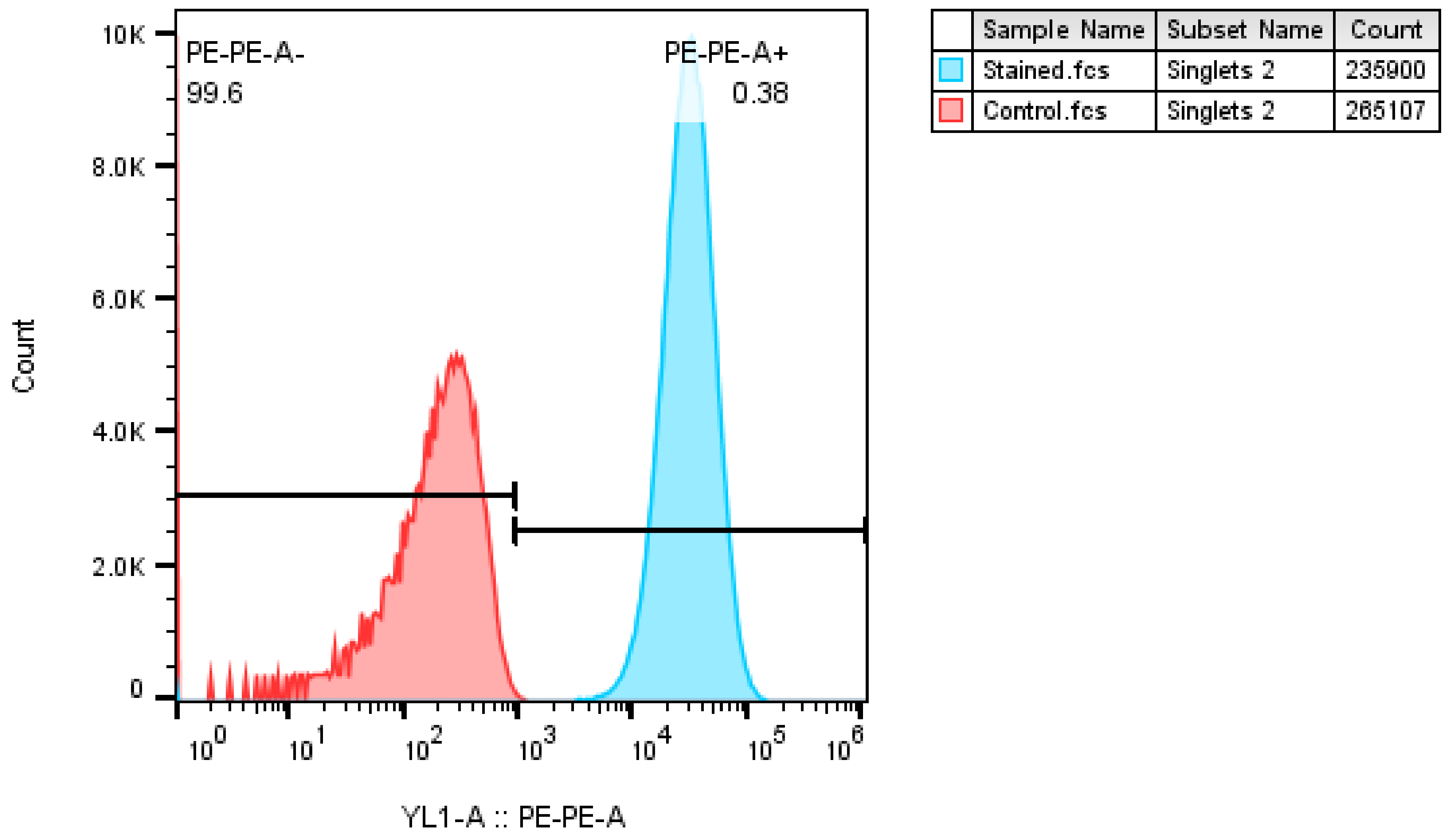


CD22

**Supplementary Figure 5. Flow data confirming the expression of CD19 and CD22 in NALM6 cells.**
NALM6 cells were stained with either APC anti-human CD19 Antibody (A) or PE anti-human CD22 Antibody (B) and subjected to flow analysis. Unstained control cells were used to determine background fluorescence.
