## Supplementary Table 1 for "Circular single-stranded DNA is a superior homology-directed repair donor template for efficient genome engineering"

**Supplementary Table 1. sgRNA and primer sequences used in this study**

| **sgRNA** | **Sequence (5' to 3')** |
|---|---|
| *RAB11A* sgRNA | *GGTAGTCGTACTCGTCGTCG* |
| *B2M* sgRNA | *GGCCACGGAGCGAGACATCT* |
| *TRAC* sgRNA | *AGAGTCTCTCAGCTGGTACA* |
| *AAVS1* sgRNA | *GGGGCCACTAGGGACAGGAT* |

| **Primer/Probe** | **Sequence (5' to 3')** |
|---|---|
| cssDNA Forward | TGGGCCATCGCCCTGATAGA |
| cssDNA Reverse | AGAATAGCCCGAGATAGGGTTGAGT |
| cssDNA probe | TCTTTAATAGTGGACTCTTGTTCCAAACT |
| dsDNA Forward | GCTTAATCAGTGAGGCACCTATC |
| dsDNA Reverse | GCCCTCCCGTATCGTAGTTAT |
| dsDNA probe | CGTTCATCCATAGTTGCCTGACTCCC |
